# Bulked segregant analysis RNA-seq (BSR-Seq) validated a stem resistance locus in *Aegilops umbellulata*, a wild relative of wheat

**DOI:** 10.1101/599597

**Authors:** Erena A. Edae, Matthew N. Rouse

**Affiliations:** Department of Plant Pathology, University of Minnesota, St. Paul, MN 55108; USDA-ARS, Cereal Disease Laboratory, St. Paul, MN 55108

**Keywords:** RNA-Seq, Stem rust resistance, GBS, *Aegilops umbellulata*, wild relatives

## Abstract

Many disease resistance genes that have been transferred from wild relatives to cultivated wheat have played a significant role in wheat production worldwide. *Ae. umbellulata* is one of the species within the genus *Aegilops* that have been successfully used as sources of resistance genes to leaf rust, stem rust and powdery mildew. The objectives of the current work was to validate the map position of a major QTL that confers resistance to the stem rust pathogen races Ug99 (TTKSK) and TTTTF with an independent bi-parental mapping population and to refine the QTL region with a bulk segregant analysis approach. Two F_2_ bi-parental mapping populations were developed from stem rust resistant *Ae. umbellulata* accessions (PI 298905 and PI 5422375) and stem rust susceptible accessions (PI 542369 and PI 554395). Firstly, one of the two populations was used to map the chromosome location of the resistance gene. Later on, the 2^nd^ population was used to validate the chromosome location in combination with a bulk segregant analysis approach. For the bulk segregant analysis, RNA was extracted from a bulk of leaf tissues of 12 homozygous resistant F3 families, and a separate bulk of 11 susceptible homozygous F_3_ families derived from the PI 5422375 and PI 554395 cross. The RNA samples of the two bulks and the two parents were sequenced for SNPs identification. Stem rust resistance QTL was validated on chromosome 2U of *Ae. umbellulata* in the same region in both populations. With bulk segregant analysis, the QTL position was delimited within 3.2 Mbp. Although there were a large number of genes in the orthologous region of the detected QTL on chromosome 2D of *Ae. tauschii*, we detected only two *Ae. umbellulata* NLR genes which can be considered as a potential candidate genes.

## Introduction

Wheat (*Triticum aestivum* L.) is used as a main source of protein and starch for human consumption. However, its production is threatened by many diseases including stem rust caused by *Puccinia graminis* f. sp. *tritici* (*Pgt*), which is known to cause severe yield losses in susceptible cultivars. Ug99, the stem rust race group discovered in Uganda in 1998, was found to be virulent on widely deployed resistance genes, and the majority of the world’s wheat germplasm is susceptible to this race group [1]. The emergence and continued evolution of Ug99 has pressed the research community to evaluate available genetic resources for resistance and rapidly develop Ug99-resistant cultivars. Since the cultivated wheat gene pool has a narrow genetic base for resistance to Ug99 and up to 90% of world’s wheat cultivars are considered Ug99 susceptible [2], identifying resistance genes from wild relatives and introgressing them into cultivated wheat is a tractable strategy for improving disease resistance.

Many wild relatives of cultivated wheats have been used as sources of disease resistance genes over the last century, and those deployed genes have also played a significant role in wheat production worldwide [3]. Among wild relatives of wheat, *Aegilops umbellulata* has been successfully used as a source of resistance genes to leaf rust, stem rust and powdery mildew [4–6]. Most recently, a major QTL conferred resistance to the *Pgt* races TTKSK (Ug99) and TTTTF was mapped on chromosome 2U of *Ae. umbellulata* using genotyping-by-sequencing (GBS) SNP markers [7].

RNA-sequencing is an efficient and cost-effective method of identifying SNPs in transcribed genomic regions especially for non-model species with a little or no available genomic resources [8–10]. The recent development of methods of SNPs detection in the transcriptome allowed the combination of bulk segregant analysis (BSA) [11] and sequencing to fine map genic regions for traits of interest. This process, known as bulked segregant analysis RNA-seq (BSR-Seq), has been used for identifying candidate genes and gene cloning purpose for cereals such as maize [12, 13] and hexaploid wheat [14, 15]. BSR-Seq method has also the ability to identify differentially expressed genes (DFGs). However, BSR-Seq may not be effective for complex traits controlled by many minor genes or traits influenced by environmental factors [16]. Most recently, a major QTL conferred resistance to the *Pgt* races TTKSK (Ug99) and TTTTF was mapped on chromosome 2U of *Ae. umbellulata* using genotyping-by-sequencing (GBS) SNP markers [7]. As part of the validation and refined mapping of this previously detected stem rust resistance QTL in *Ae. umbellulata*, two approaches were followed in the current work; independent mapping in another bi-parental population and BSR-Seq. Therefore, the objectives of the current work are to: 1) Confirm the position of previously detected major QTL from *Ae. umbellulata* 2) map the QTL identified on chromosome 2U to a shorted genetic region 3) identify candidate resistance genes, and 4) to generate the transcriptome assembly for *Ae. umbellulata.*

## Materials and Methods

### Genetic materials

Two F_2_ *Ae. umbellulata* populations were developed from crossing of resistant and susceptible accessions. The 1^st^ population comprised 140 F_2_ individuals that were obtained from crossing stem rust (race TRTTF) resistant accession PI 298905 and susceptible accession PI 542369. Similarly, the 2^nd^ population consisted of a total of 154 F_2_ individuals derived from a cross made between accessions PI 5422375 (resistant to race TTTTF) and PI 554395 (susceptible to race TTTTF). F_3_ families of both populations were also screened with race TTKSK from the Ug99 stem rust race group in a Biosafety level 3 (BSL-3) facility.

### Phenotyping mapping populations and QTL mapping

Both F_2_ individuals and F_3_ families of both populations were assessed for reaction to both races TTTTF (isolate 01MN84A-1-2) and TTKSK (04KEN156/04). Experimental procedures for inoculation, incubation, and disease assessment were conducted according to previously described methods [17]. Stem rust seedling infection types were scored on the 0-4 scale [18], and plants with infection types 0-2 were considered resistant whereas plants with infection types 3-4 were considered susceptible.

QTL mapping was carried out for the two populations described above. QTL mapping was facilitated by GBS SNP markers identified from sequence data from the two populations following the same procedures mentioned in [7]. In our previous work QTL identification was done only for the 1^st^ population using GBS SNPs. In the current work, however, the QTL mapping was carried out with GBS SNPs common between the two populations using composite interval mapping (CIM) in the RQTL package. Both F_2_ individuals and F_3_ families that were derived from F_2_s of both mapping populations were used for QTL analysis. To compare the similarity of the QTL mapped with the two populations, we inferred the QTL region from the GBS marker-based consensus map previously developed from these two populations [19].

### Bulk segregant samples, RNA-Seq library preparation and RNA sequencing

The 2^nd^ population (derived from PI 5422375 and PI 554395 cross) was used for bulk segregant analysis. F_3_ families were grown to differentiate heterozygous from homozygous F_2_ plants. A total of twenty plants from each F_3_ family were assessed at the seedling stage with *Pgt* isolates 01MN84A-1-2 (TTTTF) according to previously described methods (Rouse et al. 2011). Families were classified as homozygous resistant, heterozygotes, and homozygous susceptible based on response to the *Pgt* isolates tested. For bulk segregant analysis (BSA), 12 and 11 homozygous F_3_ families were selected to make resistant bulk and susceptible bulks based on their response to race TTTTF, respectively.

Total RNA was extracted from leaf tissues collected from homozygous resistant and susceptible families and the two parents using the RNeasy Plant Mini Kit (QIAGEN) according to the manufacture’s instruction. At least 500ng RNA of four samples (two bulks and the two parents PI 5422375 and PI 554395) was submitted to the University of Minnesota Genomics Center (Saint Paul, MN) for quality assessment, standard cDNA library preparation and RNA sequencing on Illumina HiSeq 2500 to generate 125 bp paired-end (PE) sequence reads for all four samples. Libraries were created using dual indexed TruSeq-stranded RNA library preparation kits. All libraries were pooled and sequenced across 3 lanes. The pooled libraries were Caliper XT size-selected to have inserts of approximately 200 bp. The reads were concatenated across lanes, and a single fastq file per sample for each read was generated. The average quality scores were above Q30 for all libraries.

### *De novo* transcript assembly, SNPs identification and bulk frequency ratio (BFR) calculation

Sequence read quality was evaluated using the FastQC program (http://www.bioinformatics.babraham.ac.uk/projects/fastqc/). The Trimmomatic program [20] was used to clean the reads and remove the Illumina paired-end adaptor sequences requiring an average quality score of 15, and a minimum length of reads of 30 base-pairs. Only paired-end reads that passed the cleaning steps were used for the subsequent analysis. We used Trinity assembly program [21] for *de novo* assembling the reads from the two parents. SNPs identification was done using two approaches: *de novo* assembly of the parents as a pseudo-reference, and a reference-based approach using the *Ae. tauschii* reference sequence [22].

In the *de novo* assembly approach, SNPs were identified from both the two parental assemblies (PI 5422375 and PI 554395 accessions) using the sort, merge, and pileup programs in combination with VarScan software [23]. We used the susceptible parent assembly as a pseudo reference to call SNPs from short reads of the pools. Fisher exact tests with a 0.05 significance level were applied to identify SNPs that showed frequency differences between the two pools. However, only SNPs that were common between the parental SNPs and the two pools were considered as true SNPs. Only reads with base phred quality scores of 20 or more, and requiring a minimum number of 6 reads were considered for SNPs for both pools. Then the parental SNPs were identified from the bulks based on SNP positions by merging the parental SNPs with SNPs identified from the pools. Additionally, bulk frequency ratio was calculated based on the allele frequency of the resistant allele in the pools to identify SNPs-linked to stem rust resistance. Bulk frequency ratio (BFR) is defined as the allele frequency ratio of the resistant allele (resistant parent allele) in the resistant bulk compared to the resistant allele frequency in the susceptible bulk. BFR ratio of four was used as the minimum threshold to identify potential candidate SNPs.

### Reference-based identification of SNPs using *Ae. tauschii* as a reference sequence

SNP calling was also completed using *Ae. tauschii* v4.0 as a reference (Luo et al. 2017). SNP calling was done for the bulks and the parents separately. For the two bulks, the short reads were aligned against the *Ae. tauschii* reference sequence using BWA and the SNPs were called using samtools [24] and Varscan softwares [23]. SNP quality filtering parameters were set at; minimum read depth at a position to make a call 10, minimum supporting reads at a position to call variants 6, minimum average base quality for variant-supporting reads 20, and average variant frequency of 0.2. The default was used for the remaining parameters. Similarly, to call SNPs from the two parents, the RNA-Seq short reads of both parents were aligned against the *Ae. tauschii* v4.0 reference sequence using BWA and SNPs were called with samtools software with default set. VCFtools was used to filter out false SNPs with the following filtering criteria: minor allele frequency of >0.20, phered-quality score of 20, minimum read depth of 6 and average mapping quality (MQ) of 30.

### Sequence assembly characterization and functional annotation

Both *de novo* assemblies of *Ae. umbellulata* accessions were aligned against local genomic and cDNA databases of hexaploid wheat, *Ae. tauschii* and barley. Transcripts were also annotated using BLAST homology searches against the UniProt databases within the Trinotate functional annotation site (https://trinotate.github.io/). Briefly, the likely coding regions were predicted using Transcoder software and then the predicted proteins were searched in the protein database by BLAST. In addition, protein domains were searched using “hmmer” software [25]. The predicted proteins and protein domain search results were integrated into coding region selection. Finally, all search results were uploaded to “sqlite” database to extract annotation reports with Trinotate suite.

We also assessed if there were genes that contained a nucleotide-binding site and leucine-rich repeat (NLR) domain in the QTL region. For this purpose, all contigs from the resistant parent assembly that were mapped on chromosome 2D of *Ae. tauschii* through blast approach were used to identify contigs potentially containing NLR domains in this region. Motif Alignment and Search Tool (MAST) in MEME suite [26] and NLR-parser, a java program that identifies NLRs from MAST motifs output [27], were used to identify contigs that comprised NLR motifs using a set 20 previously characterized amino acid sequences that contain NB-LRR domains [28], and six-frame translated amino acid sequences derived from contig sequences that were mapped on chromosome 2D of *Ae. tauschii.*

## Results

### QTL mapping using two bi-parental mapping populations

GBS markers common across the two mapping populations were used to identify stem rust resistance QTL. Detailed procedures of GBS marker development have been reported previously for these bi-parental populations (Edae et al. 2017). QTL mapping was done separately for each population. A major stem rust resistance QTL was detected on chromosome 2U of *Ae. umbellulata* at the same region with both mapping populations (Fig. 1). However, QTL interval size in the 1^st^ population (derived from accession PI 298905 and PI 542369 cross) is larger (5.86 cM based on 1.5-lod interval and 0.95 Bayes confidence interval analyses) than that of the 2^nd^ population (accession PI 5422375 and PI 554395 cross) which was estimated to be 1.68 cM. GBS marker TP34301 (consensus position: 87.89 cM) that was linked with the QTL in the 1^st^ population was only 1.35 cM away from another GBS marker TP44300 (consensus position: 89.24 cM) linked with the QTL (based on race TTTTF) in the 2^nd^ population, and both markers are shown in bold face in Fig. 2. The markers flanking the entire region of the QTL (83.7 - 89.5 cM) are colored red. Based on the two populations, the QTL is bounded by GBS markers TP17463 (83.7 cM) and TP24305 (89. 5 cM) within a 5.8 cM interval. Thus, the target for fine mapping with BSR-Seq approach is this 5.8 cM QTL region.

**Fig. 1.**
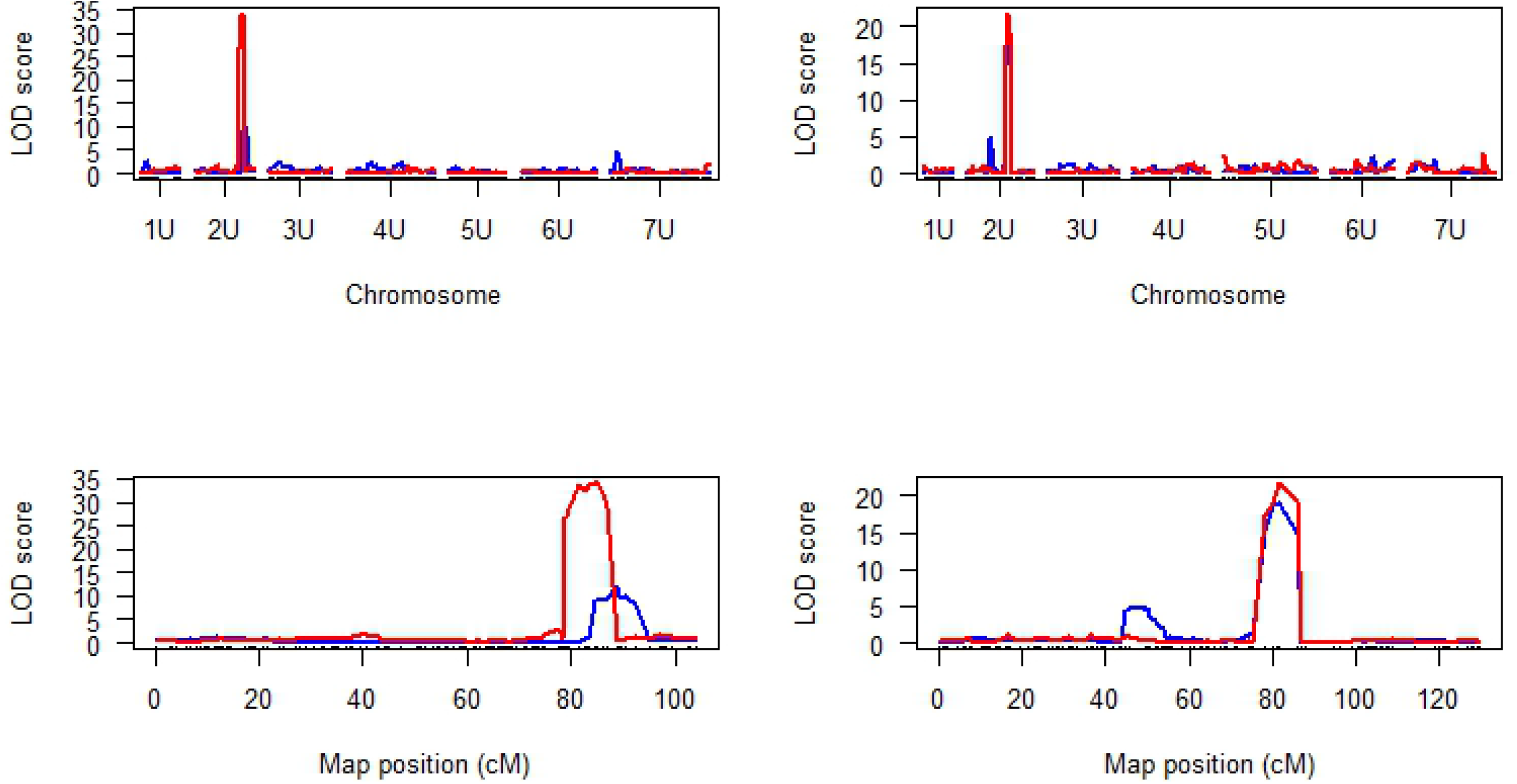
Chromosome and logarithm of odds (LOD) profile of stem resistance QTL detected for races TTTTF and TTKSK in two the *Ae. umbellulata* F_2:3_ populations. The 1^st^ population is on the left picture whereas the 2^nd^ population is indicated on the right picture.

**Fig. 2.**
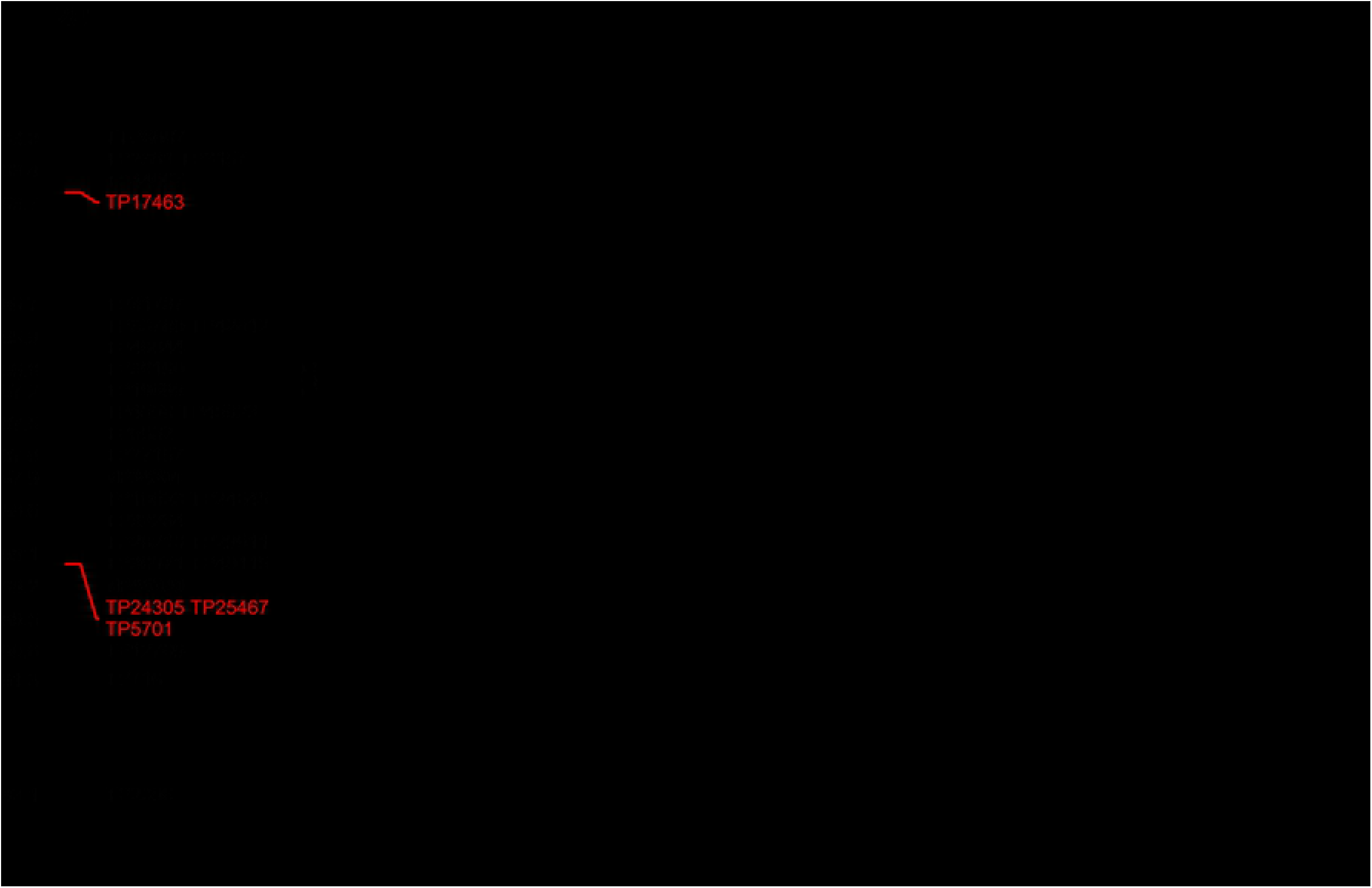
Stem resistance QTL region mapped from two *Ae. umbellulata* bi-parental populations.

The SNP context sequences of all GBS markers were aligned against the pseudomolecule of the *Ae. tauschii* reference, and all markers linked with resistance against *Pgt* races TTTTF and TTKSK in both populations were within 7.89 Mbp (566677717- 574572533 bp) on chromosome 2D of *Ae. tauschii* (Fig. 3).

**Fig. 3.**
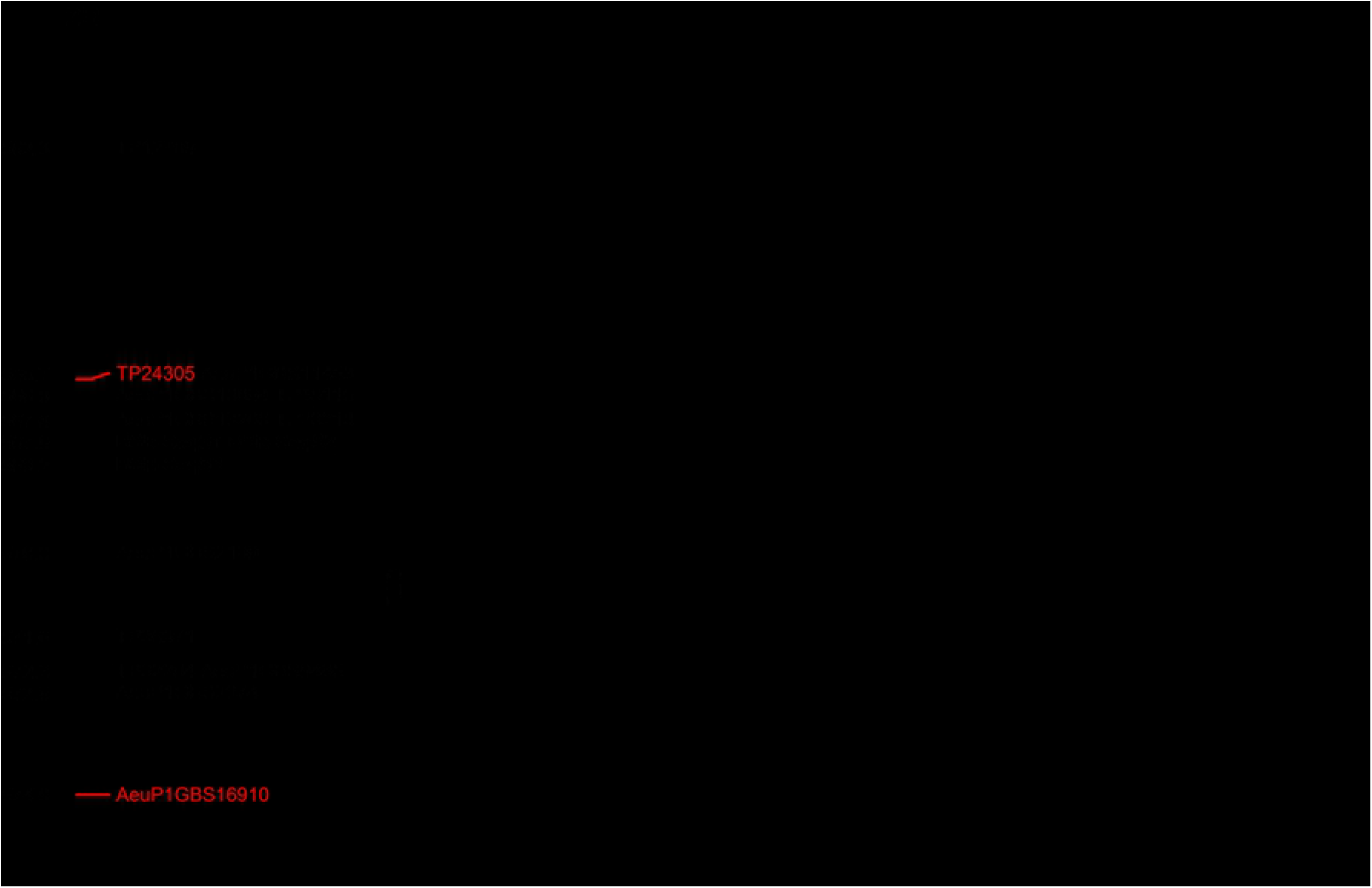
Physical map position of the stem resistance QTL based on *Ae. tauschii* reference sequence.

### Transcriptome assemblies of *Ae umbellulata*

The number of contigs were 201,588 and 239,727 for the *de novo* assembly of resistant and susceptible parents, respectively (Table 1). After clustering with cd-hit program [29], the contig number was reduced to 164,456 and 194,020 for the resistant and susceptible accessions with N50 values 1,548 and 1,829, respectively.

**Table 1.** Summary of basics statistics of the primary assemblies generated for PI 5422375 and PI 554395 genotypes of *Ae. umbellulata*.

| Genotype | Contigs | Contig mean size<br>(bp) | N50 | GC% |
| --- | --- | --- | --- | --- |
| PI 5422375 | 201,588 | 889.5 | 1,548 | 48.17 |
| PI 554395 | 239,727 | 1061.50 | 1,829 | 47.58 |

**Table 2.**
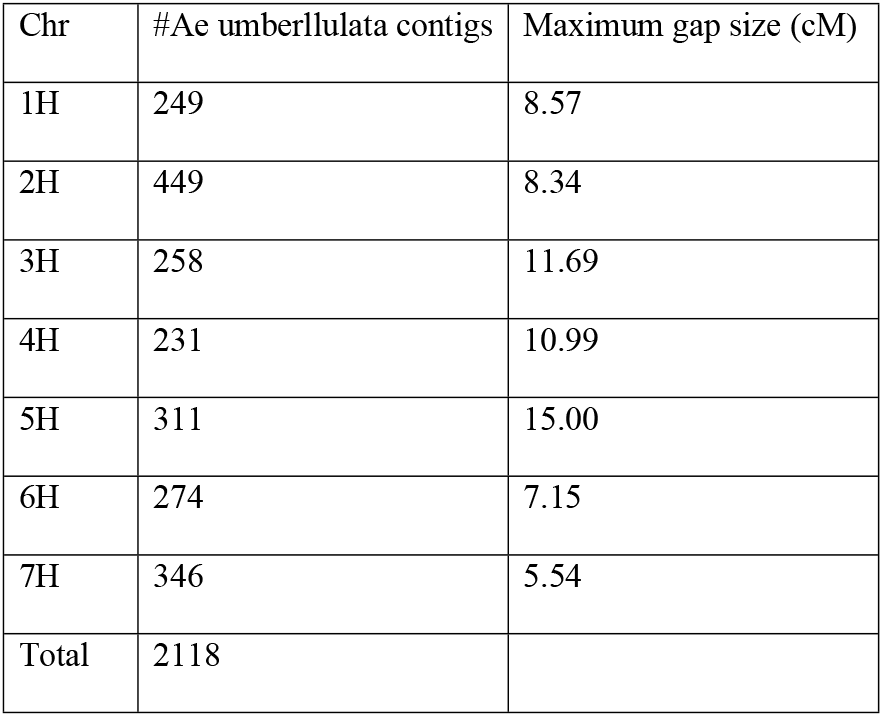
Distribution of SNP-containing contigs from susceptible *Ae. umbellulata* accession PI 554395 assembly across barley chromosomes.

| Chr | #*Ae. umbellulata* contigs | Maximum gap size (cM) |
| --- | --- | --- |
| 1H | 249 | 8.57 |
| 2H | 449 | 8.34 |
| 3H | 258 | 11.69 |
| 4H | 231 | 10.99 |
| 5H | 311 | 15.00 |
| 6H | 274 | 7.15 |
| 7H | 346 | 5.54 |
| Total | 2118 |  |

**Table 3.** QTL detected for resistance to two *Pgt* races on chromosome 2U with two populations of *Ae umbellulata* with composite interval (CIM) and multiple QTL mapping methods (MQM).

| Linked marker | Consensus position (CM) | Method | population | LOD | Race |
| --- | --- | --- | --- | --- | --- |
| TP34301 | 87.89 | CIM | 1<sup>st</sup> population | 30.8 | TTTTF |
| TP44300 | 89.24 | CIM | 2<sup>nd</sup> population | 21.78 | TTTTF |
| TP44300* | 89.24 | CIM | 2nd population | 19.16 | TTKSK |
| TP5701 | 89.47 | MQM | 1st population | 17.78 | TTTTF |
| TP5701 | 89.47 | MQM | 1st population | 5.9 | TTKSK |
| TP27197 | 87.56 | MQM | 2nd population | 24.78 | TTKSK |

\*QTL flanking markers

### *Aegilops tauschii* as a reference for SNP identification

Using *Ae. tauschii* as a reference genome sequence, we identified a total of 400,327 SNPs, and the highest number of SNPs was recorded on chromosome 2D (correspond to 2U of *Ae. umbellulata)* whereas the lowest number of SNPs was found on chromosome 4D (Fig. 4). Out of these, three SNPs within the QTL region (566.7-574.6 Mbp) showed large allele frequency differences between the two bulks. These three SNPs were located within 0.3 Mbp and two of them were within 14 bp. Most importantly, one of these three SNPs (position 567,944,494 bp) had the largest bulk frequency ratio (BFR) of 97.08 of all SNPs in the dataset, and another SNP (position 567,944,392 bp) was also ranked 3^rd^ with BFR value of 89.29 (Table S1). These two SNPs are very close (0.39 Mbp) to the GBS markers TP13266 and TP28218 that were linked to stem rust resistance QTL in the 1^st^ and the 2^nd^ mapping population, respectively.

**Fig. 4.**
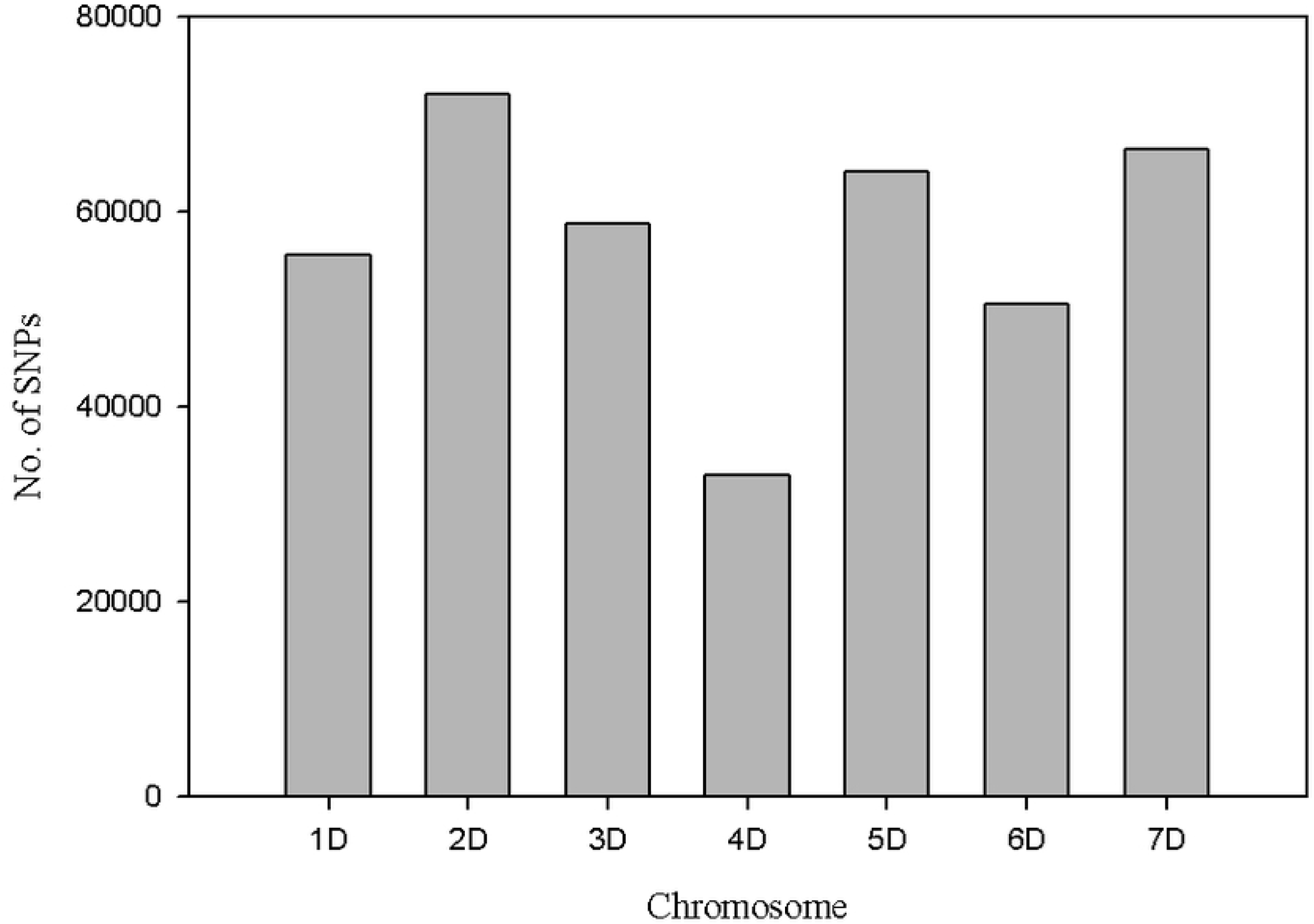
Distribution of SNPs identified from *Ae. umbellulata* assembly on *Ae. tauschii* chromosomes.

A total of 1,237 SNPs with BFR greater or equal to four with no heterozygotes in the two bulks were obtained. Out of these, 81% (1003) SNPs were on chromosome 2D of *Ae. tauschii* and the remaining 234 SNPs were distributed on the remaining six chromosomes (Fig. 5). Moreover, out of 129 SNPs with BFR >20, a total of 95% (123) SNPs were on chromosome 2D. Only 4.6% (6) SNPs were false positive. Among the SNPs on chromosome 2D, a majority with high BFR values mapped in the QTL region detected with the bi-parental QTL mapping approach (Fig. 6). Therefore, we predicted, based on GBS QTL mapping and bulk segregant analysis, the stem rust resistance QTL identified with *Ae. umbellulata* bi-parental mapping populations is in the orthologous region of 566.68-569.96 Mbp on the physical map of chromosome 2D of *Ae. tauschii.* The size of this refined region is approximately 3.2 Mbp. All genes in this region were extracted from the *Ae. tauschii* transcribed region, and there were a total of 111 predicted genes in the region (Table S2).

**Fig. 5.**
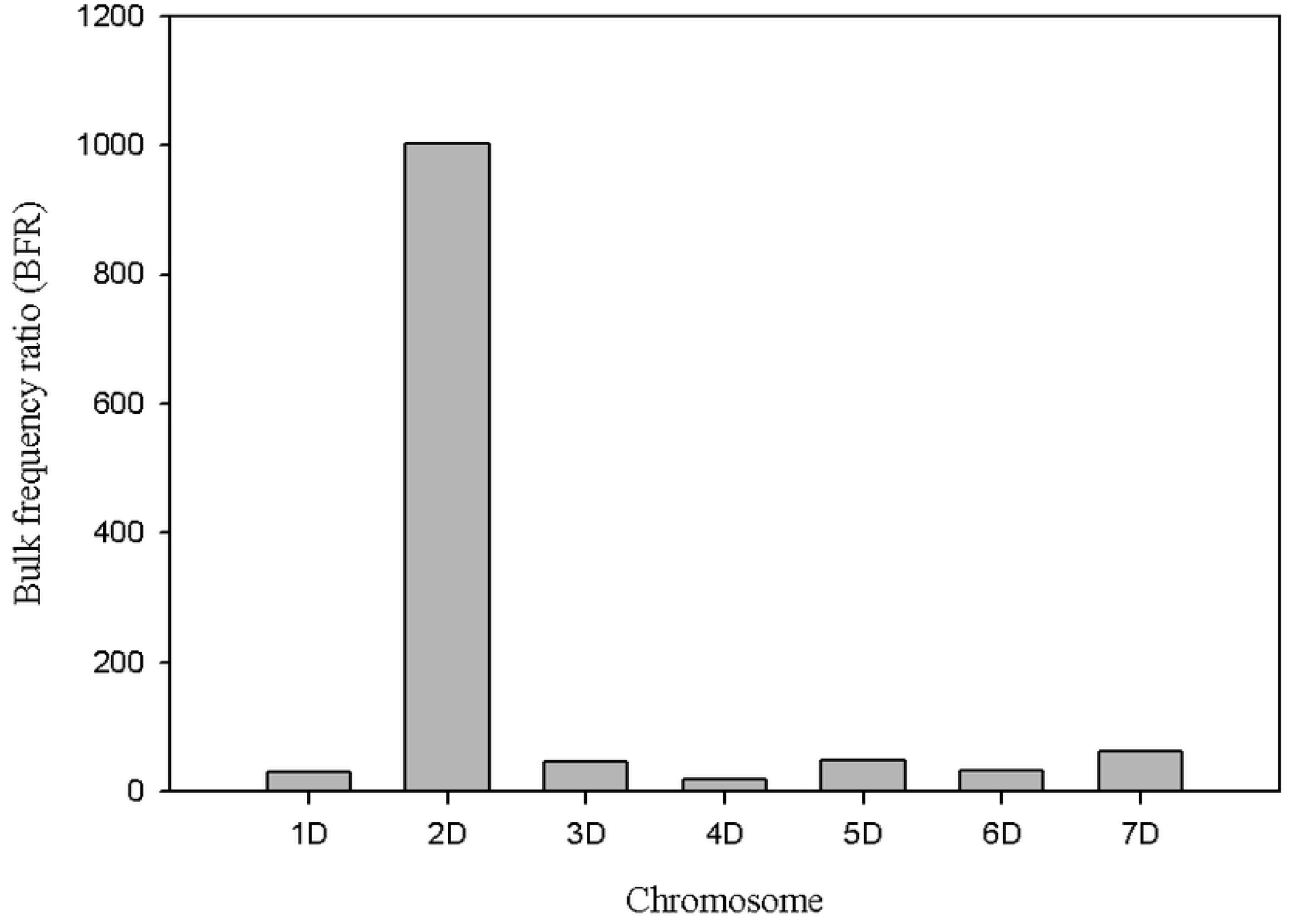
Distribution of SNPs with bulk frequency ratio higher than 4 on *Ae. tauschii* chromosomes.

**Fig. 6.**
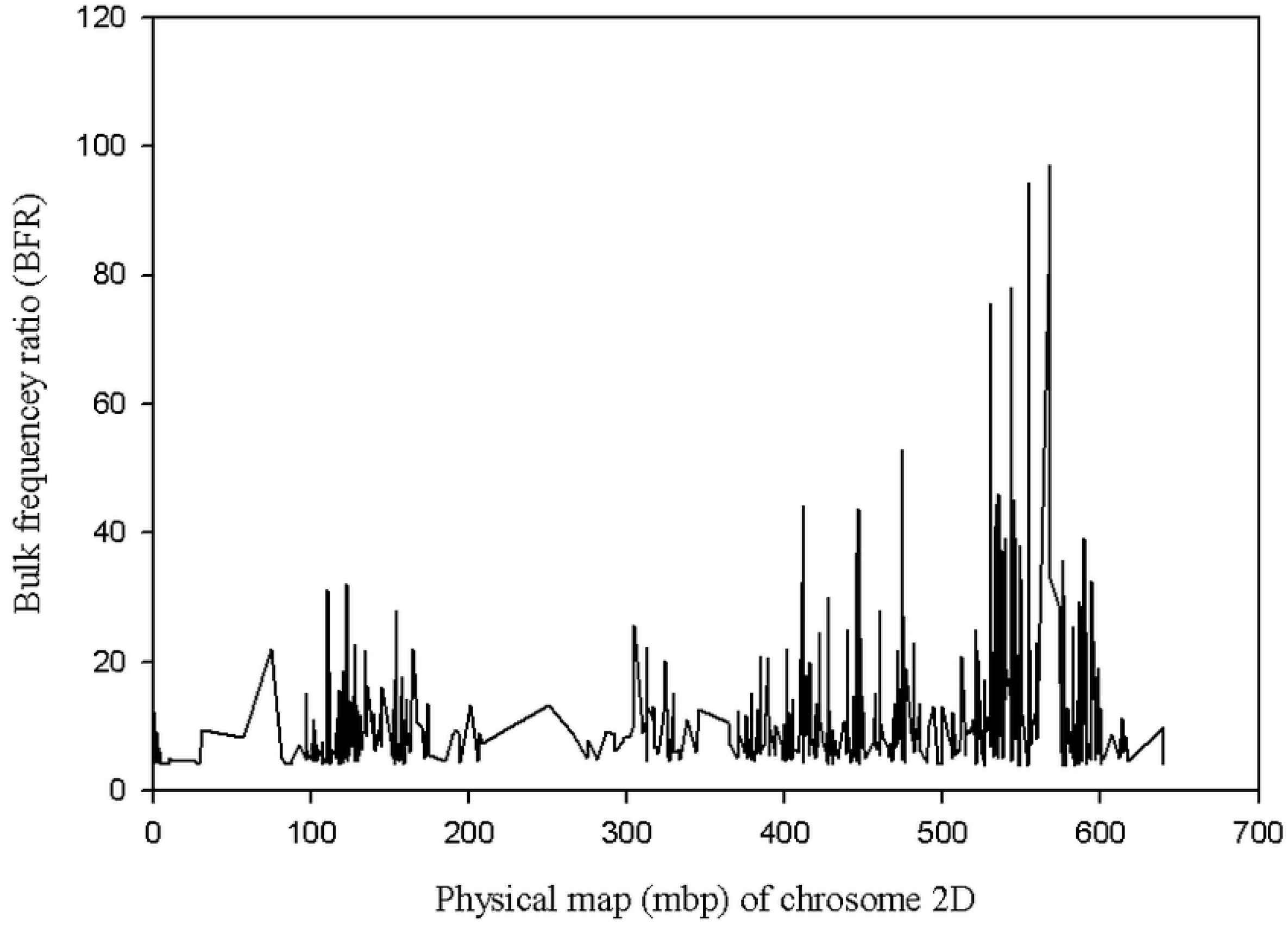
Distribution of SNPs with bulk frequency ratio higher than 4 on chromosome 2D *Ae. tauschii* chromosome.

For the purpose of assessing the presence of genes that contain a nucleotide-binding site and leucine-rich repeat (NLR) domain in the QTL region, contigs of the resistant parent assembly that mapped on chromosome 2D of *Ae. tauschii* were used. Among 34,906 resistant contigs that had a significant blast hit on chromosome 2D, a total of 147 contigs contain an NLR domain (partial domains included). Out of these, only four contigs are in the predicted gene region, of which DN39311_c0_g2_i1, DN39311_c0_g2_i4 and DN39311_c0_g2_i5 are isoforms of one gene and mapped on the same physical position (567.98 Mbp) on chromosome 2D. The remaining NLR contig, DN16173_c0_g1_i1, was mapped 0.94 Mbp away from the former (at 568.93 Mbp on chromosome 2D of *Ae. tauschii* physical map). Interestingly, from BSR-Seq analysis, two SNPs with the highest BFR (97.08 and 89.29) for the two bulks (resistant and susceptible) were identified at 567.94 Mbp on chromosome 2D suggesting that these three isoforms represent potential candidate genes for the QTL that we mapped with the bi-parental populations (Fig. 7; Table S3). The similarity searches in the NCBI database for one of the three contigs (DN39311_c0_g2_i1, DN39311_c0_g2_i2, DN39311_c0_g2_i4 and N39311_c0_g2_i5) retrieved CDS that encoded NBS-LRR disease resistance protein in barley (rga S-127 gene) and hexaploid wheat (NBS-LRR resistance protein CIN14 mRNA) as top hits. Similarly, the identity search for DN16173_c0_g1_i1 retrieved *Brachypodium distachyon* disease resistance protein RGA2-like (LOC100839373) but none of the hits was NBS-LRR protein.

**Fig. 7.**
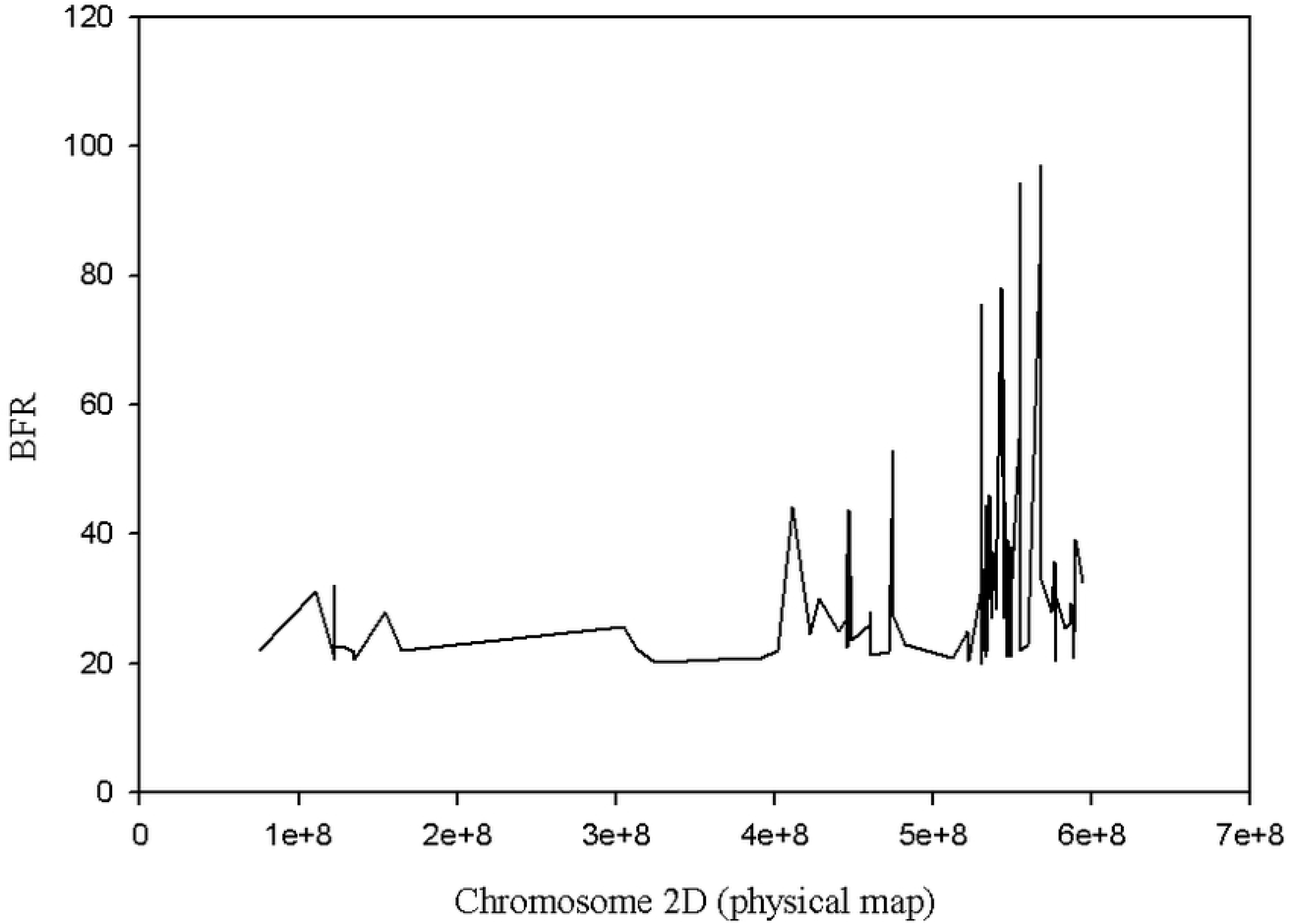
Distribution of SNPs with NLR domain on chromosome 2D of *Ae. tauschii* chromosome.

### *De novo* SNPs identification

Using the susceptible parent assembly as a reference, a total of 103,970 unique SNPs were identified with default settings of samtools and VarScan softwares. Out of these, 99,170 SNPs had significant best hits in the *Ae. tauschii* reference sequence and 98.8% (97,987) of the SNPs had known physical positions. The remaining 1% mapped to unmapped scaffolds. Chromosome 2U had the highest number of SNPs whereas chromosome 4U had the least number of SNPs, and the number of markers were between the two extremes for the remaining six chromosomes (Fig. 8). The blast similarity search of the susceptible parent assembly in the *Ae. tauschii* local data base retrieved hits for a total of 203,271 contigs. Out of these, a total of 201,065 had known physical map positions and the remaining contigs mapped to unmapped scaffolds. In terms of distribution of contigs across chromosomes, although the lowest number of contigs was for chromosome 4D, generally the seven chromosomes can be grouped broadly in two. Chromosomes 2D, 3D, 5D and 7D each compose greater numbers of contigs (29,000-33,000) whereas chromosomes 1D, 4D and 6D possess lower numbers of contigs (23,000-26,000) (Fig. 9).

**Fig. 8.**
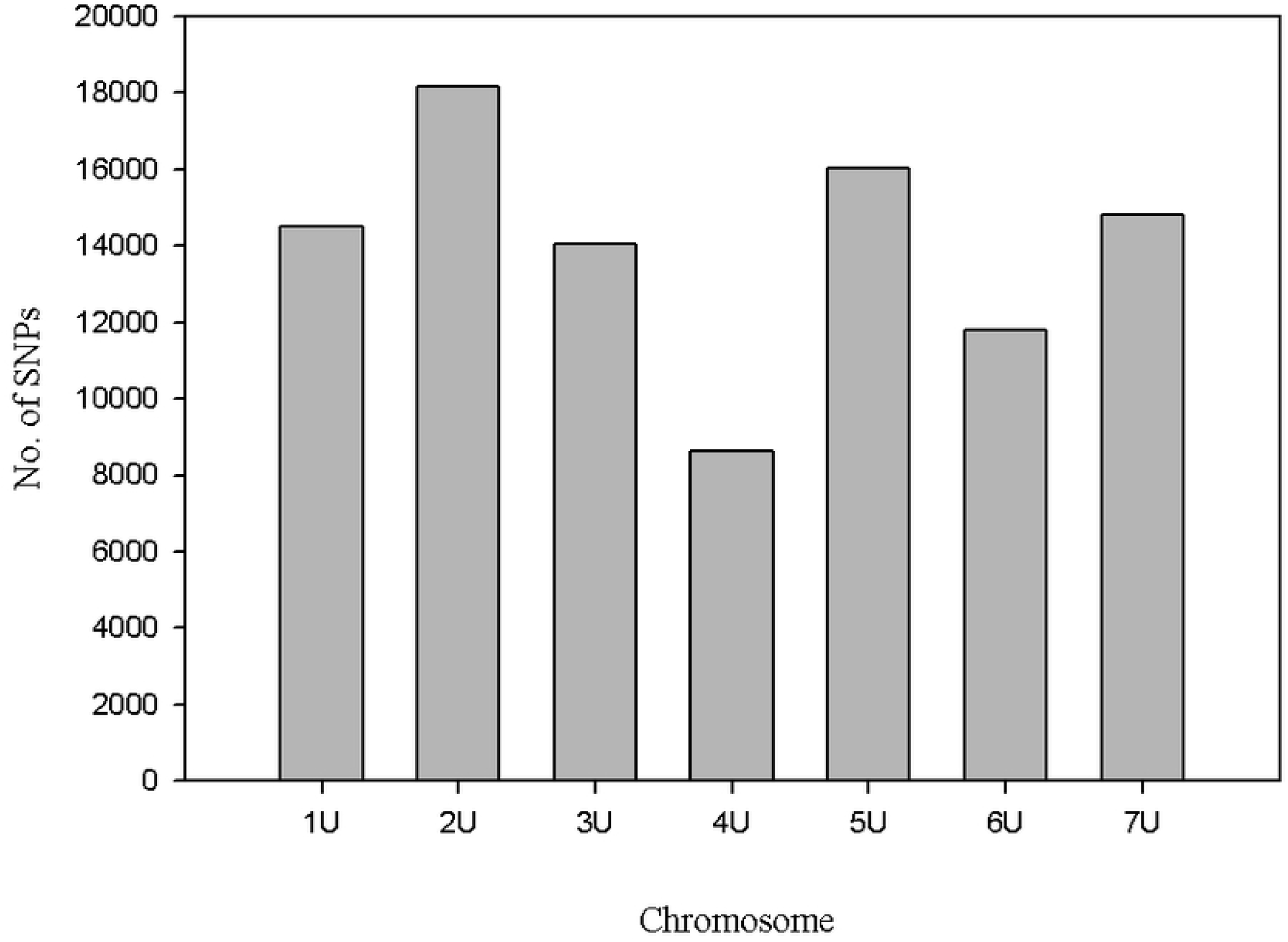
Distribution of SNPs identified from susceptible parent’s assembly with *de novo* approach on *Ae tauschii* chromosomes.

**Fig. 9.**
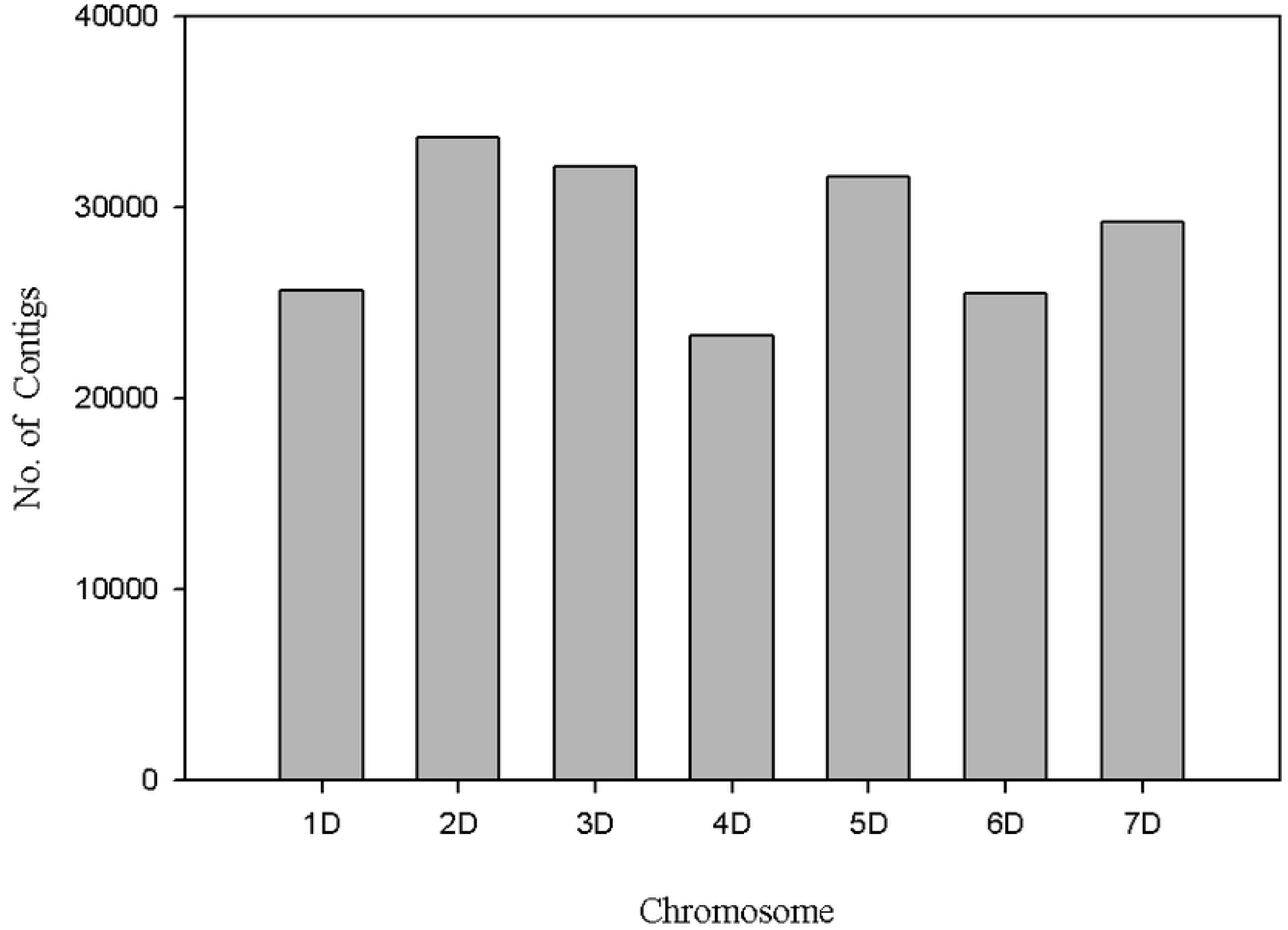
Distribution of contigs in the susceptible parent’s assembly on the *Ae. tauschii* chromosomes.

SNPs were also called for the two pools using the *de novo* assembly approach. A total of 25,931 unique SNPs were obtained with the default setting. In order to identify the parental SNPs in the two pools, we merged the parental SNP dataset with that of the bulks’ SNP data sets using the parental SNP positions to link the two SNP data sets. A total of 12,311 parental SNPs were identified from the bulks and merged with output from blasting the susceptible parent assembly with the *Ae. tauschii* reference sequence. Of these, a total of 11, 764 had significant hits in the reference sequence of *Ae. tauschii.*

### Identification of SNPs associated with stem rust resistance using *de novo* assembly

The susceptible parent assembly was aligned against the *Ae. tauchii* genome reference sequence, and a total of 36, 973 unique contigs had hits on chromosome 2D, and then these unique contigs were again merged with the 25, 931 SNPs identified using the susceptible parent as a pseudo reference. Out of these, a total of 6,346 SNPs were from chromosome 2D, of these, 2,222 SNPs had BFR values of four and above (Table S4). The number of SNPs with BFR values of 20 and above was 394. There were a total of three SNPs that are scored both in parents and bulks with BFR values of 4 and above in the identified QTL region (562.3-574.6 Mbp on chr 2D). The SNPs within the contig DN19592_c0_g1_i showed the highest BFR values and mapped within the QTL region 0.01 Mbp (at position 562305371 on chr 2D) away from TP12079 that was linked with the QTL. There were a total of nine SNPs in this contig with high BFR values. The 2^nd^ SNP DN19324_c0_g1_i2 (position on chromosome 2D: 566490221 pb) with BFR value of 119 was mapped at position 566.49 which is also in the QTL region. The 3^rd^ contig DN2794_c0_g1_i1 (574564921 bp) with a SNP associated with stem rust resistance had a BFR value of 6.77.

### Clustering the *Ae umbellulata* assemblies into genes

To predict the number of genes in *Ae. umbellulata*, the two assemblies were aligned against the high confidence gene sets of hexaploid wheat, *Ae. tauschii* and barley. Contigs from *Ae umbellulta* that shared a high confidence CDS with hexaploid wheat were considered as isoforms of a gene. Out of the total 204,037 contigs from the resistant parent assembly, 59% (120,819) transcripts had hits in the hexaploid wheat gene model IWGSC RefSeq Annotation v1.0. These hits were grouped into the gene model identification numbers they matched. The resistant parent hits matched a total 43,276 genes in the 137,914 high confidence genes of hexaploid wheat. Out of these, a total of 42,807 genes had known chromosome information in the hexaploid wheat gene annotation file. Similarly, out of 239,727 transcripts in the susceptible parent assembly, 52% (125,293) transcripts were assigned into 44,027 unique high confidence genes in hexaploid wheat. The database comprised a total of 137,056 high confidence genes of hexaploid wheat. The similarity search in the 39, 622 high confidence gene set of *Ae. tauschii* retrieved a total of 22,685 and 23,111 unique genes for resistant and susceptible accession assemblies. Similarly, integrating the resistant parent into the 26,159 high confidence gene model of barley, *Ae. umbellulata* transcripts matched a total of 18,669 unique genes.

### Annotation of transcriptome

For the two assemblies, 40% and 39% of all transcripts matched records in Uniprot for the resistant parent and susceptible parents, respectively (Table S5). Gene Ontology (GO) terms were assigned to transcripts matching GO-annotated records in the Uniprot database to characterize the biological functions from sequence similarity. The distribution of the GO terms across the three domains were highly comparable for the two assemblies (Supplemental Fig. 1).

## Discussion

Wild relatives of wheat have been used as sources of resistance genes in wheat improvement programs for many decades. *Ae. umbellulata* is one of those wild species that has been a useful source of resistance genes. Leaf rust resistance *Lr9* was the 1^st^ gene that was transferred to wheat 60 years ago from *Ae. umbellulata*, the secondary gene pool of wheat [30]. *Ae. umbellulata* has also been identified as a source of resistance to stem rust [31, 32, 7], powdery mildew, Hessian fly and green bug [4]. In the current work a new stem rust resistance gene from *Ae. umbellulata* was mapped on chromosome 2U, and its map position was validated with independent mapping bi-population, and finally the region was fine-mapped via integration of bi-parental QTL mapping method with the bulk segregant RNA-Seq analysis (BSR-Seq) approach.

BSR-Seq has been recently applied for fine-mapping gene region in cereal crops such as wheat [14, 15]and maize [12, 13]. It is a powerful method to enrich gene region with SNPs derived only from expressed portion of a genome thereby facilitates identification the actual gene itself. Using SNP markers that showed large allele frequency difference between resistance and susceptible bulks, the stem resistant QTL region was validated and refined with the BSR-Seq approach in this report.

We used comparative mapping to infer the map positions of QTL-linked GBS markers and RNA-Seq derived SNPs using recently released *Ae tauschii* reference sequence v.1.0, and *de novo* assemblies of *Ae. umbellulata.* From the integration of the SNP context sequences from RNA-Seq and GBS data into a pseudomolecule of *Ae. tauschii*, the QTL-associated GBS markers were mapped in the same region with the RNA-Seq SNPs that showed the highest BFR ratio, suggesting the suitability of BSR-Seq for validating and identifying candidate genes detected originally as QTL in classical mapping. Once the gene region was confidently known, by combining information from the physical positions of all GBS markers that were associated with the resistance gene in two bi-parental populations and RNA-Seq SNPs with high BFR) the size of gene interval was approximately 3.2 Mbp based on physical map of chromosome 2D of *Ae. tauschii.* Given stem rust resistance genes are known to encode nucleotide-binding site and leucine rich repeat (NLR) proteins [33–35], and the QTL identified from *Ae. umbellulata* bi- parental populations fits into features of a qualitative gene, exploring the presence of NLR genes in the predicted region is useful to identify short lists of potential candidate genes. In this way, five contigs with NLR domains were discovered from the resistant parent assembly that had hits within this 3.2 Mbp gene region, of which four NLR contigs are variants of a single gene. The similarity search of the four variants of this gene indicated that the gene encodes disease resistance NLR proteins in barley and hexaploid wheat suggesting the NLR contigs represent a potential candidate gene for the major stem resistance QTL we mapped on chromosome 2U of *Ae umbelluata.* Moreover, this result agrees with our previous hypothesis that the gene identified from *Ae umbellulata* is a novel gene as none of the candidate NLR contigs had similarity to the known major disease resistance genes in barley and hexaploid wheat.

Two approaches were followed for the identification of SNPs that showed differences between the resistant and susceptible bulks to fine map the QTL region: *de novo* method and orthologous reference-based approach. Although SNPs were effectively identified with the *de novo* approach where *de novo* assemblies were used as a pseudo-references to call SNPs from RNA-Seq data, orthologous reference sequence-based approach enabled us to precisely identify SNPs that are closer to the region of the gene mapped with GBS markers with two bi-parental mapping populations. This indicated that with the absence of a species reference sequence the use of closely related species reference sequence is an advantageous alternative for SNPs identification. SNPs with high BFR from both methods mapped at the same region, but the resolution was better in the cross-species reference-based approach as we identified more SNPs with higher BFR that mapped closer to the GBS marker position.

In summary, our study demonstrated that BSR-Seq is cheap and efficient approach not only precisely validating the map position of previously identified QTL but also identification of potential candidate genes is possible as the approach provides additional SNPs in the mapping interval. BSR-Seq could also be utilized for other wild relatives of crop species that are difficult to propagate and self-fertilize.

---

Supplemental Fig. 1. Distribution of GO (Gene Ontology) terms across categories for resistant parent assembly.

